# Non-Parametric Genetic Prediction of Complex Traits with Latent Dirichlet Process Regression Models

**DOI:** 10.1101/149609

**Authors:** Ping Zeng, Xiang Zhou

**Affiliations:** Department of Epidemiology and Biostatistics, Xuzhou Medical University, Xuzhou, Jiangsu 221004, China; Department of Biostatistics, University of Michigan, Ann Arbor, Michigan 48109, USA.; Center for Statistical Genetics, University of Michigan, Ann Arbor, Michigan 48109, USA.

**Author notes:** Correspondence and requests for materials should be addressed to XZ.

## Abstract

Using genotype data to perform accurate genetic prediction of complex traits can facilitate genomic selection in animal and plant breeding programs, and can aid in the development of personalized medicine in humans. Because most complex traits have a polygenic architecture, accurate genetic prediction often requires modeling all genetic variants together via polygenic methods. Here, we develop such a polygenic method, which we refer to as the latent Dirichlet process regression model (DPR). DPR is non-parametric in nature, relies on the Dirichlet process to flexibly and adaptively model the effect size distribution, and thus enjoys robust prediction performance across a broad spectrum of genetic architectures. We compare DPR with several commonly used prediction methods with simulations. We further apply DPR to predict gene expressions, to conduct PrediXcan based gene set test, to perform genomic selection of four traits in two species, and to predict eight complex traits in a human cohort.

## Introduction

Genome-wide association studies (GWASs) have identified thousands of genetic loci harboring associated single nucleotide polymorphisms (SNPs) for many complex traits and diseases, providing unprecedented insights into the genetic basis of phenotypic variation^1-8^. The accumulation of genetic data from existing association studies has led to a growing interest in predicting traits and diseases using genetic markers (in addition to using traditional environmental or clinical variables)^9^. In animals or plants, accurate phenotype prediction with genetic markers can assist the selection of individuals with desirable breeding values and can improve the effectiveness of breeding programs^10^. In humans, accurate phenotype prediction with genetic markers can facilitate disease prevention and intervention at early stages and can aid in the development of personalized medicine by using genotype information to customize the treatment and predict the outcome^11^. Phenotype prediction has also been proposed recently as a key step for integrating functional genomic sequencing studies with GWASs: we can construct more powerful and interpretable gene-set tests in GWASs by setting variant weights to be the coefficients inferred from predictive models in expression quantitative trait locus (eQTL) mapping studies^12^.

Progress towards accurate phenotype prediction requires the development of statistical methods that can model all SNPs jointly. Previous association studies have demonstrated that most complex traits and common diseases have a polygenic background and are each influenced by many genetic variants with small effects. For instance, it is estimated that thousands of causal variants influence human height^13^. Similarly, many animal or plant traits are contributed by hundreds of causal variants (e.g. maize-related traits, such as kernel oil and growing degree days^14,15^; and cattle-related traits, such as backfat thickness, milk yield and hot carcass weight^16,17^). Because most complex traits and common diseases have a polygenic architecture, a handful of identified associated SNPs often only capture a small proportion of the phenotypic variation and thus cannot be used to yield accurate phenotype and risk prediction. Instead, accurate phenotype prediction requires polygenic models that can make use of all genome-wide SNPs^9,18-20^. In the past decade, successful development and application of many polygenic models in the context of genomic selection has revolutionized many animal breeding programs^16,21-23^. More recently, applications of polygenic models to human GWASs have also yielded fruitful results^11,24-27^.

Most existing polygenic models for prediction make an assumption on the effect size distribution and different methods differ mainly in such modeling assumption. For example, the commonly used linear mixed model (LMM), also known as the best linear unbiased predictor (BLUP), assumes that the effect sizes from all variants follow a normal distribution^9,28^. The Bayes alphabetic (e.g. BayesA and BayesB) methods assume that the variant effect sizes follow a t-distribution or its variation^10,18,29^. The Bayesian lasso assumes a double exponential/Laplace distribution^30,31^. NEG generalizes the Bayesian lasso by assuming a normal exponential gamma distribution^32^. BVSR and BayesCπ assume a point-normal distribution^29,33^. BSLMM assumes a mixture of two normal distributions^34^ and is closely related to the early reversible jump Markov Chain Monte Carlo (rjMCMC) method^20^. BayesR^35^ assumes a three-component normal mixture together with a point mass at zero. Given the large number of modeling choices, one naturally wonders which method to use for any given trait. Previous studies have suggested that accurate prediction requires choosing a prior effect size distribution that can closely match the shape of the true effect size distribution, such that the inferred posterior can approximate well the polygenic architecture of the given trait^24,35,36^. However, the effect size distribution for any given trait or disease is unknown *a priori* and varies for different diseases in terms of the number of causal variants, their minor allele frequencies (MAFs), and their individual effect sizes^11^. Therefore, to achieve robust performance, it is important to design prior distributions that are flexible enough to resemble the true effect distribution in many traits as close as possible^34,35^.

Up to now, almost all existing polygenic models are parametric in nature and use a prior effect size distribution that is characterized by a few parameters. From the information channel perspective^37^, the number of parameters in a parametric model determines model complexity and bounds the amount of information in data that can be captured by the model^37-40^. Therefore, using only a few parameters to characterize the effect size distribution can limit the flexibility of the model^37,38^ and impede its robust performance across a range of genetic architectures. As an example, the commonly applied LMM uses a normal distribution with one variance component parameter to characterize the effect size distribution. For highly polygenic traits, the assumed normal distribution can approximate the true effect size distribution well, and as a result, LMM can achieve good predictive performance^34,35^. However, for traits with large effect variants, the assumed normal distribution can no longer capture the true effect size distribution well and the performance of LMM decays^34,35^.

To allow for greater flexibility on the *a priori* effect size distribution and to enable robust phenotype prediction performance across a range of phenotypes, we develop a Bayesian non-parametric model, which we refer to as the latent Dirichlet process regression (DPR). DPR does not use any fixed parametric distribution as the prior choice for the effect size distribution. Instead, DPR relies on the Dirichlet process to assign a prior on the effect size distribution itself and is thus capable of inferring an effect size distribution from the data at hand. Effectively, DPR uses infinitely many parameters *a priori* to character the effect size distribution, and with such a flexible modeling assumption, DPR is capable of adapting to a broad spectrum of genetic architectures and achieves robust predictive performance across a wide range of complex traits. We illustrate the benefits of DPR with simulations and real data applications for gene expression prediction, gene-based test via PrediXcan, genomic selection for four traits in two species, as well as genetic prediction of eight complex traits in a human cohort.

## Results

### Method overview

An overview of our method is provided in the Methods section with details provided in the Supplementary Note. Briefly, we use a Dirichlet process to introduce a non-parametric effect size distribution that can robustly resemble a large classes of unimodal distributions. Indeed, our prior effect size distribution can be used to adaptively and accurately approximate a t-distribution, a point-t mixture distribution, a mixture of step functions, as well as the marginal effect sizes estimated from a real data set; whereas a normal distribution cannot (Fig. 1). Therefore, our prior distribution on the effect size can adaptively approximate a wide range of possible effect size distributions underlying complex traits. Since accurate modeling of the effect size distribution is a key to achieve accurate prediction performance^24,34,36^, we expect our non-parametric model to perform robustly well across a range of polygenic architectures. Our method is implemented in the DPR software, freely available at http://www.xzlab.org/software.html.

**Figure 1.**
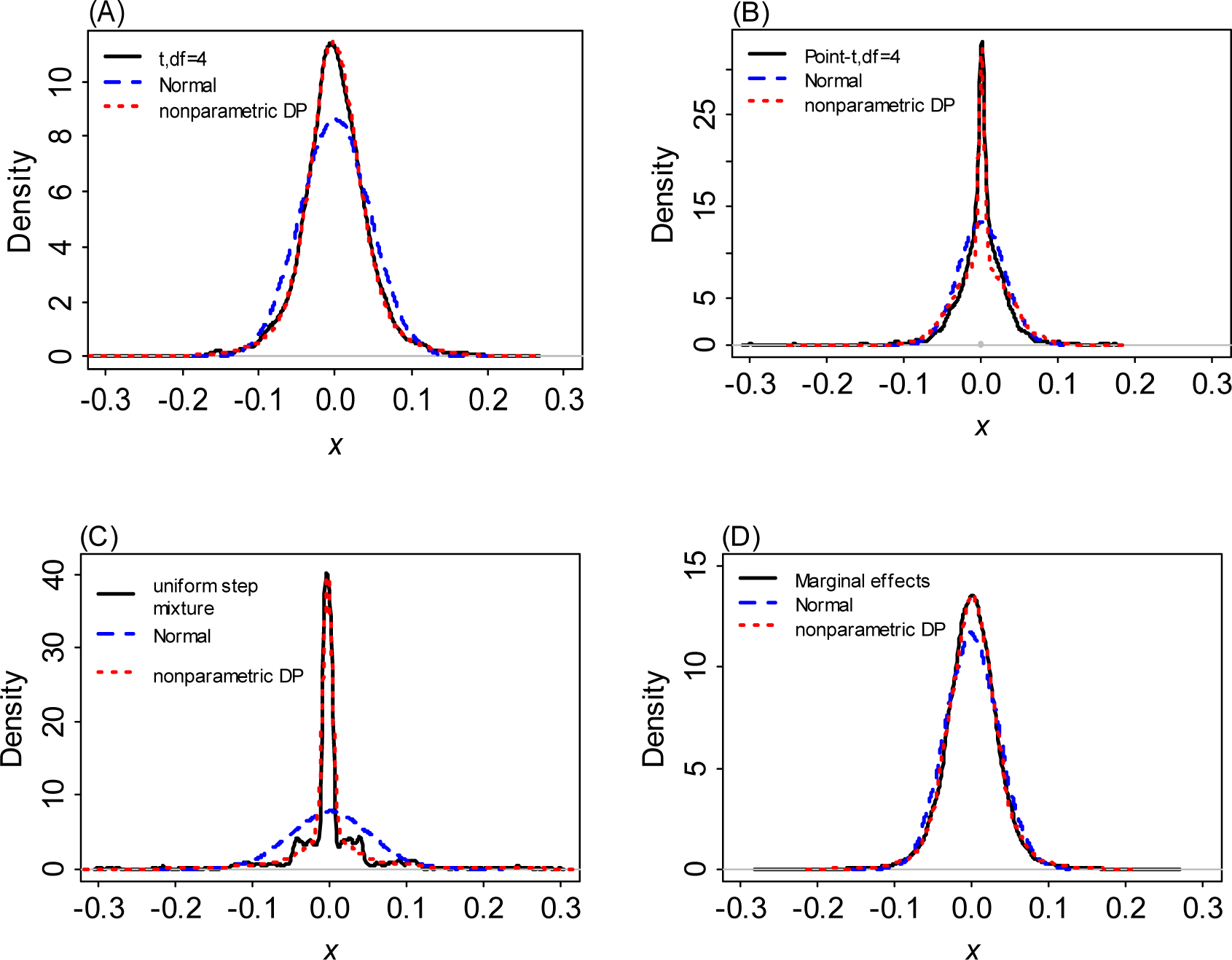
The induced non-parametric Dirichlet process (nonparametric DP) normal mixture prior on the effect sizes can be used to approximate a large number of unimodal distributions. We either simulated 2,000 values from (A) a standard t-distribution with df=4; (B) a point-t mixture distribution with the zero proportion being 0.2, or equivalently, 0.8×t(df=4)+0.2×*δ*_0_, where 0 denotes a point mass at zero; (C) a four-component uniform step mixture distribution 0.50×U(-0.05,0.05)+0.25×U(- 0.3,0.3)+0.15×U(-0.8,0.8)+0.05×U(-2,2), where U denotes a uniform distribution; or obtained (D) the estimated marginal effect sizes from a linear mixed model in the cattle data with SCS (somatic cell score) as the phenotype. To make the first three data comparable with the last data in (D), we centered and scaled the values from the first three data for them to have a mean of zero and within the range of (-0.3,0.3). We then fit each data with either our non-parametric distribution (red) or a normal distribution (blue), and displayed the fitted curves on top of the sample distribution (black). Clearly, the non-parametric Dirichlet process normal mixture can approximate all these distributions well, while a simple normal distribution cannot.

### Simulations

We first compare the performance of DPR with several other commonly used prediction methods using simulations. A total of seven different methods are included for comparison: (1) BVSR^29^; (2) BayesR^35^; (3) LMM^28^; (4) MultiBLUP^41^; (5) rjMCMC^20^; (6) DPR.VB, the variational Bayesian (VB) version of DPR; and (7) DPR.MCMC, the Markov chain Monte Carlo (MCMC) version of DPR. Note that both BayesR and MultiBLUP have been recently demonstrated to outperform a range of existing prediction methods; thus, we do not include other prediction methods into comparison here.

To make our simulations as real as possible, we used genotypes from an existing cattle GWAS dataset^17^ with 5,024 individuals and 42,551 SNPs and simulated phenotypes. To cover a range of possible genetic architectures, we consider eight simulation settings from four different simulation scenarios with the phenotypic variance explained (PVE) by all SNPs being either 0.2, 0.5, or 0.8 (details in Methods). In each setting for each PVE value, we performed 20 simulation replicates. In each replicate, we randomly split the data into a training data with 80% individuals and a test data with the remaining 20% individuals. We then fitted different methods on the training data and evaluated their prediction performance on the test data (i.e. Monte Carlo cross validation). We evaluated prediction performance using either the squared correlation coefficient (*R*^2^) or mean squared error (MSE). We contrasted the prediction performance of all other methods with that of DPR.MCMC by taking the difference of *R*^2^ or MSE between the other methods and DPR.MCMC. Therefore, an *R*^2^ difference below zero or an MSE difference above zero suggests worse performance than DPR.MCMC. Fig. 2 shows *R*^2^ differences for different methods across 20 replicates in each of the eight simulation settings for PVE=0.5. Because Fig. 2 shows prediction performance difference, a large sample variance of a method in the figure only implies that the prediction performance of the method differs a lot from that of DPR.MCMC, but does not imply that the method itself has a large variation in predictive performance. Supplementary Table 1 shows the standard deviation of absolute *R*^2^ values across cross variation replicates; various methods display similar prediction variability. Supplementary Figs 1 and 2 show the *R*^2^ differences for PVE=0.2 and PVE=0.8, respectively. The corresponding results for MSE differences are shown in Supplementary Figs 3-5. The *R*^2^ and MSE values of the baseline method, DPR.MCMC, are shown in the corresponding figure legend.

**Figure 2.**
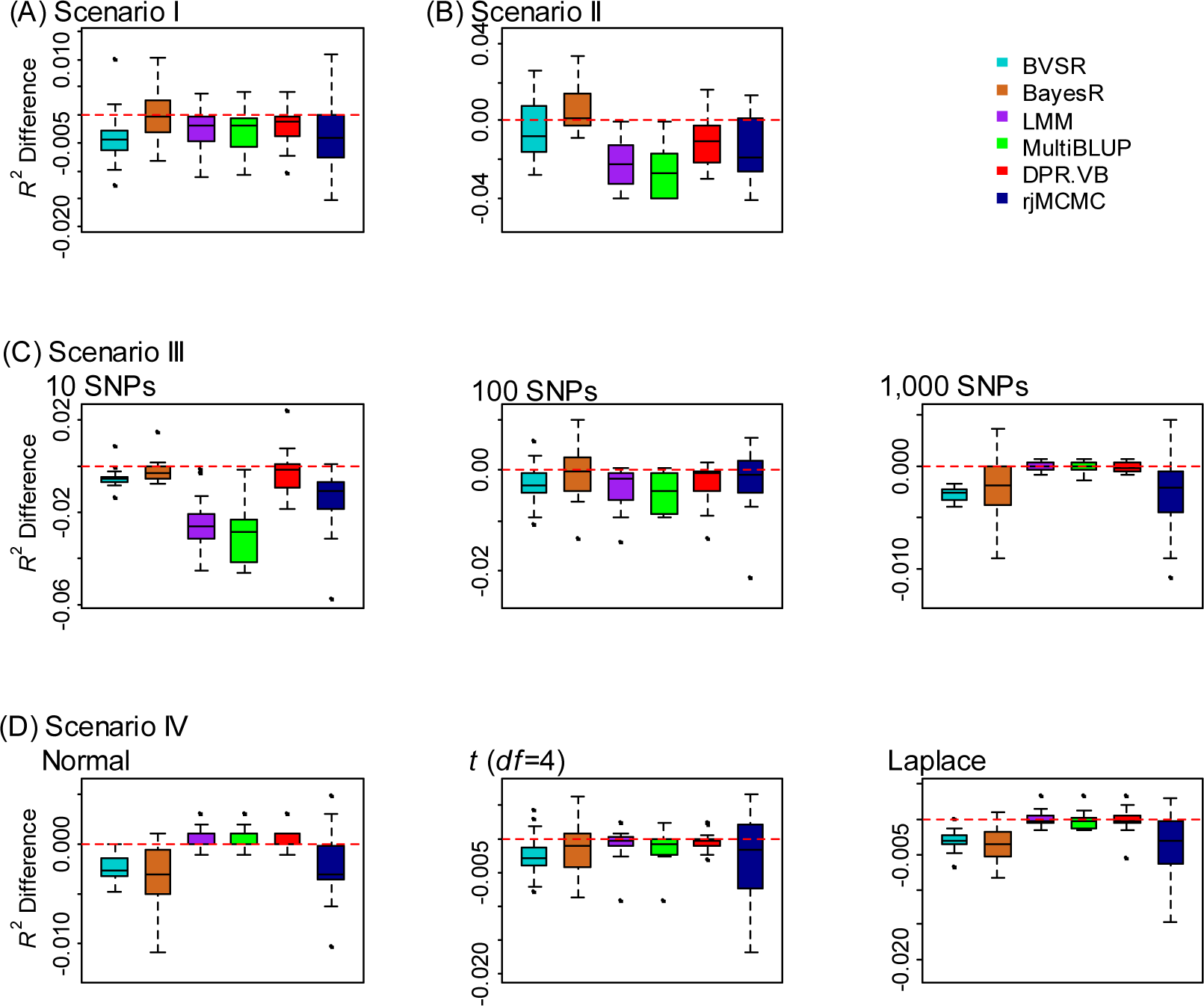
Comparison of prediction performance of several methods with DPR.MCMC in simulations when PVE=0.5. Performance is measured by *R*^2^ difference with respect to DPR.MCMC, where a negative value (i.e. values below the red horizontal line) indicates worse performance than DPR.MCMC. The sample *R*^2^ differences are obtained from 20 replicates in each scenario. Methods for comparison include BVSR (cyan), BayesR (chocolate), LMM (purple), MultiBLUP (green), DPR.VB (red), rjMCMC (black blue) and DPR.MCMC. Simulation scenarios include: (A) Scenario I, which satisfies the DPR modeling assumption; (B) Scenario II, which satisfies the BayesR modeling assumption; (C) Scenario III, where the number of SNPs in the large effect group is 10, 100, or 1,000; and (D) Scenario IV, where the effect sizes are generated from either a normal distribution, a t-distribution or a Laplace distribution. For each box plot, the bottom and top of the box are the first and third quartiles, while the ends of whiskers represent either the lowest datum within 1.5 interquartile range of the lower quartile or the highest datum within 1.5 interquartile range of the upper quartile. For DPR.MCMC, the mean predictive *R*^2^ in the test set and the standard deviation for the eight settings are respectively 0.272 (0.031), 0.299 (0.026), 0.295 (0.026), 0.281 (0.030), 0.277 (0.035), 0.278 (0.030), 0.282 (0.025) and 0.273 (0.022).

Overall, while each method works the best when their individual modeling assumption is satisfied, DPR.MCMC is robust and works well across all eight settings from four scenarios. For example, if we rank the methods based on their median performance across replicates, then when the total PVE is moderate (e.g. PVE=0.5, Fig. 2; note that for each PVE there are a total of eight simulation settings for the four scenarios), DPR.MCMC is the best or among the best in seven simulation settings (i.e. scenario I, c=10, 100 and 1,000 in scenario III, and normal, t and Laplace distributions in scenario IV; where “among the best” refers to the case when the difference between the given method and the best method is within ±0.001) and is ranked as the second best in the rest one simulation setting (i.e. scenario II). Similarly, when the total PVE is high (e.g. PVE=0.8, Supplementary Fig. 2), DPR.MCMC is the best or among the best in seven simulation settings, and it is ranked as the second best in scenario IV when the effect size follows a normal distribution. Even when DPR.MCMC is ranked as the second best method, the difference between DPR.MCMC and the best method is often small. Among the rest of the methods, LMM, MultiBLUP and rjMCMC all work well in polygenic settings (scenario I; c=1,000 in scenario III; scenario IV) but can perform poorly in sparse settings (scenario II; c=10 and c=100 in scenario III). The performance of LMM, MultiBLUP and rjMCMC in polygenic vs sparse settings presumably stems from their polygenic assumptions on the effect size distribution. In contrast, because of the sparse assumption on the effect size distribution, both BayesR and BVSR have an advantage in sparse settings (scenario II; c=10 or 100 in scenario III) but suffers in polygenic settings (c=1,000 in scenario III; scenario IV). The performance of BVSR is also generally worse than BayesR in the challenging setting when PVE is either small or moderate, presumably because of the much simpler prior assumption employed in BVSR for the non-zero effects. Finally, the VB version of DPR (i.e. DPR.VB) performs considerably less well compared with the MCMC version of DPR (i.e. DPR.MCMC), especially when PVE is high (Supplementary Fig. 2). However, DPR.VB still compares favorably with the other methods when PVE is small or moderate (Supplementary Fig. 1).

### Real data applications

To gain further insights, we compare the performance of DPR with the other methods in several real data sets to (1) predict gene expression levels using cis-SNPs; (2) conduct subsequent PrediXcan based gene set test; (3) perform genomic selection in animal studies; and (4) predict complex traits in humans.

Our first application is predicting gene expression levels using cis-SNPs in the GEUVADIS data^42^. The GEUVADIS data contains gene expression measurements on 15,810 genes and 465 individuals after quality control (Methods). These individuals have their genotypes measured in the 1000 Genomes project^43^. In the data, we first identified cis-SNPs that are within 100 kb of each gene and obtained an average of 175 cis-SNPs per gene. Then, for each gene in turn, we applied different methods to predict gene expression levels using these cis-SNPs. To measure prediction performance, we carried out 20 Monte Carlo cross validation data splits as in simulations. In each data split, we fitted methods in a training set with 80% of randomly selected individuals and evaluated method performance using *R*^2^ in the test set with the remaining 20% of individuals. In addition to the seven methods used in the simulations (i.e. LMM, BVSR, MultiBLUP, BayesR, rjMCMC, DPR.VB and DPR.MCMC), we also applied Elastic Net (ENET)^44^, which is the default method used in the original PrediXcan paper^12^. Table 1 lists the number of genes with a predictive *R*^2^ above different thresholds for different methods. The predictive *R*^2^ obtained from DPR.MCMC versus various other methods across all genes is shown as scatter plots in Supplementary Fig. 6, where each plot also lists the number of genes for which DPR.MCMC performs better and the number of genes for which DPR.MCMC performs worse.

**Table 1.**
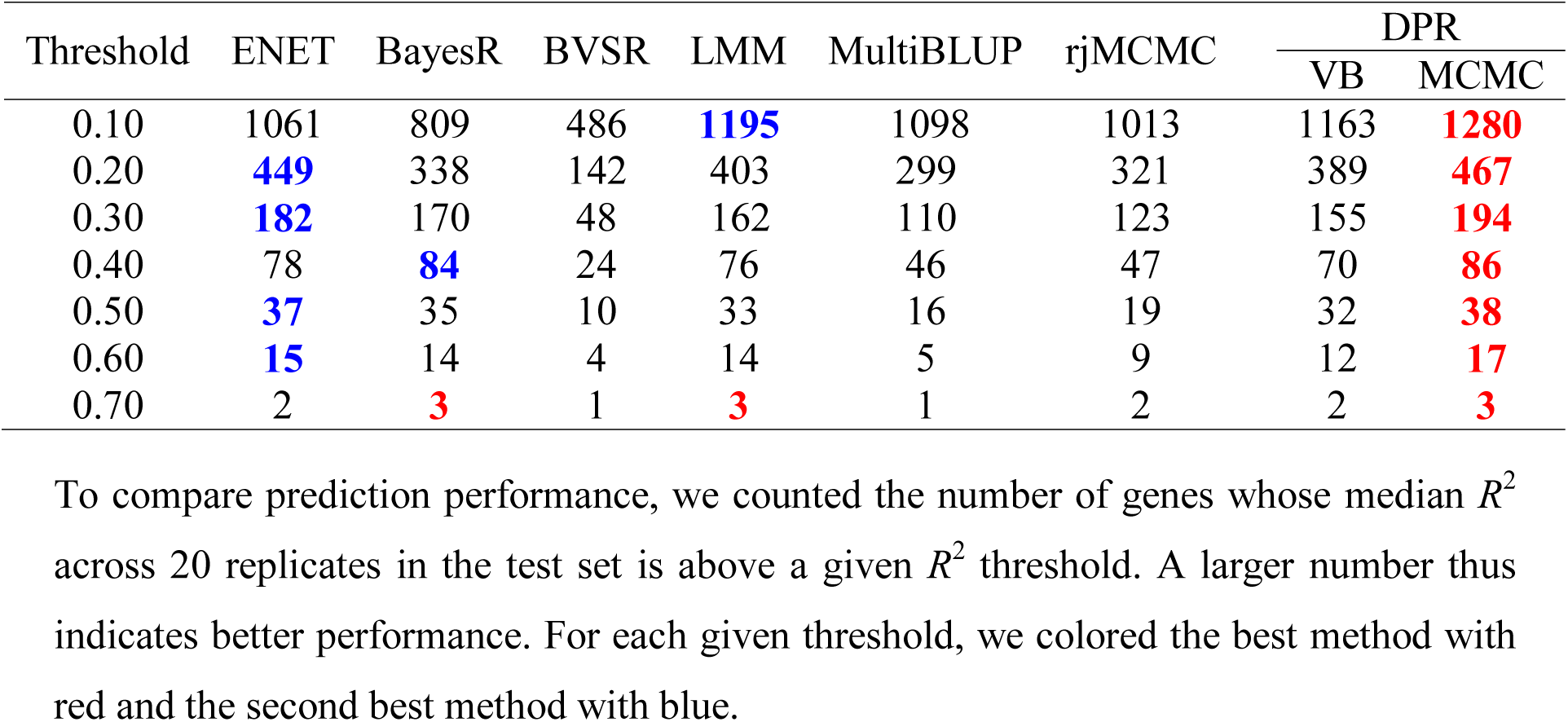
Comparison of seven different methods in predicting gene expression levels in the GEUVADIS data.

The results are largely consistent with these in simulations. Overall, DPR.MCMC generally achieves better predictive performance than the other methods. For example, DPR.MCMC is able to achieve a higher predictive *R*^2^>0.10 in ~1,300 genes, which is ~100 more than that by the second best method at this threshold (i.e. LMM; Table 1). Similarly, compared with other methods, not only does DPR.MCMC achieve a higher *R*^2^ for most genes; the *R*^2^ improvement from DPR.MCMC can be large for many genes (Supplementary Fig. 6). Among the rest of the methods, the performance of LMM, DPR.VB and ENET are comparable with each other and are ranked right behind DPR.MCMC. On the other hand, the two sparse models (i.e. BVSR and BayesR) perform poorly in this data, especially for some genes whose expression levels are highly predictive by the other methods (Table 1, Supplementary Fig. 6).

The robust performance of DPR.MCMC in predicting gene expression levels also translates to a relatively high power in the subsequent PrediXcan gene set test. To perform PrediXcan gene set test, we consider the seven common diseases from WTCCC^4^ as in Gamazon *et al*.^12^. For each disease and each gene in turn, we used the estimated cis-SNP effect sizes on expression levels from GEUVADIS as weights to construct gene set tests in WTCCC. Following Gamazon *et al*.^12^, we focused on a set of 4,343 genes that had a predictive *R*^2^ above 0.01 from all methods. The results are shown in Table 2, which lists the number of significant genes identified by different methods for different diseases. In total, DPR.MCMC identified 38 genes associated with different phenotypes, more than that identified by any other methods. The performance of DPR.MCMC is followed by DPR.VB and subsequently LMM and rjMCMC. Supplementary Table 2 lists the significant genes identified by DPR.MCMC, which are all consistent with previous studies. We also note that, in general, a higher gene expression predictive performance leads to a higher power in the subsequent gene set analysis. In addition, consistent with their relatively poor gene expression prediction performance, the two sparse models (BayesR and BVSR) do not perform well in the gene set test as well.

**Table 2.**
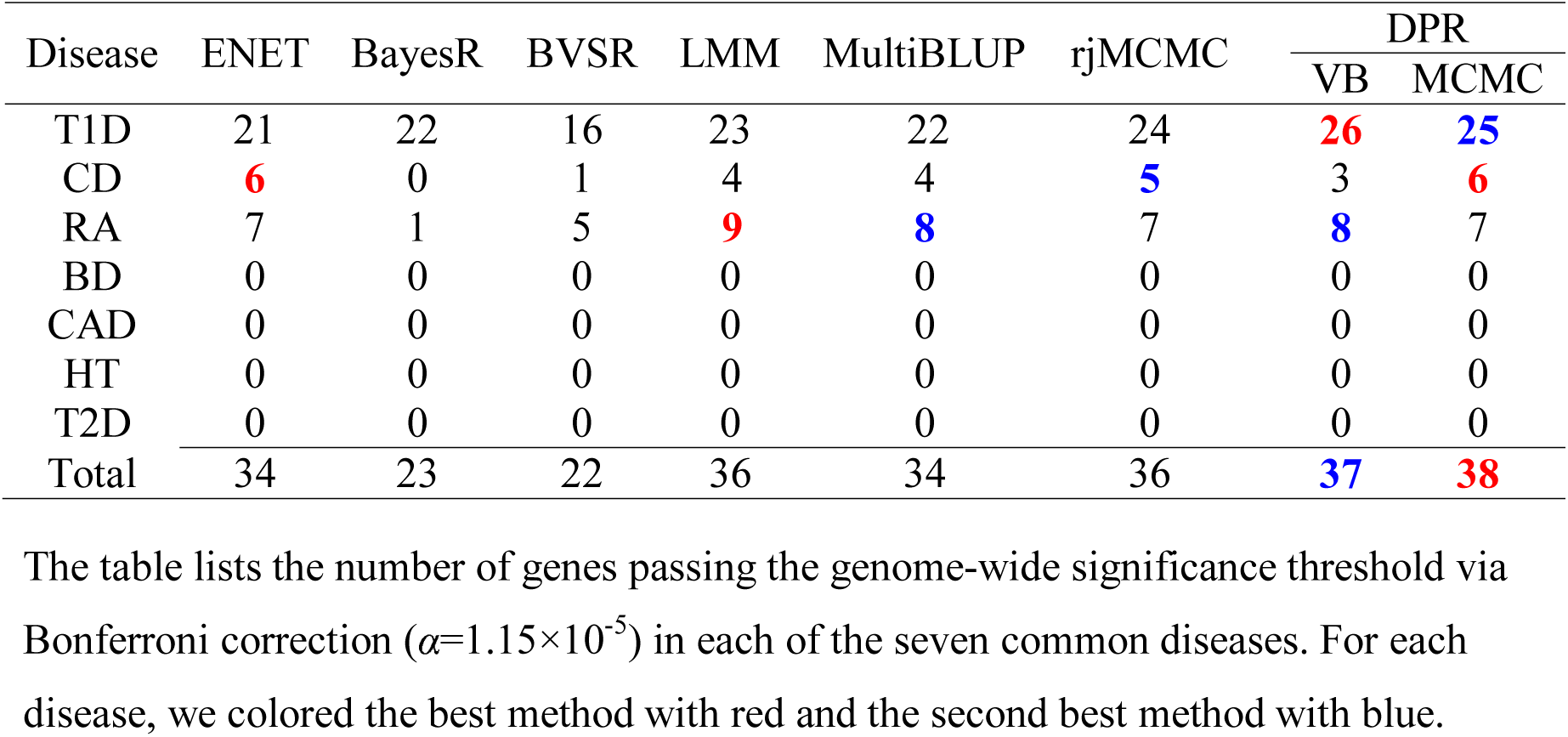
Comparison of seven different methods in the PrediXcan gene set test in the WTCCC data.

Finally, we compare the performance of DPR with the other methods in predicting phenotypes in three GWAS data sets: (1) a cattle study^17^, where we focus on three phenotypes: milk fat percentage (MFP), milk yield (MY), as well as somatic cell score (SCS); (2) a maize study^15^, where we use growing degree day (GDD) as the phenotype; (3) the Framingham heart study (FHS) data^45^, where we focus on five plasma traits that include low-density lipoprotein (LDL) cholesterol, glucose (GLU), high-density lipoprotein (HDL) cholesterol, total cholesterol (TC) and triglycerides (TG), and three anthropometric traits that include height, weight and body mass index (BMI). As in simulations, for each phenotype, we performed 20 Monte Carlo cross validation data splits. In each data split, we fitted methods in a training set with 80% of randomly selected individuals and evaluated method performance using *R*^2^ or MSE in a test set with the remaining 20% of individuals. We again contrasted the performance of the other methods with that of DPR.MCMC by taking the *R*^2^ difference or MSE difference with respect to DPR.MCMC. The results are shown in Fig. 3 (*R*^2^ difference) and Supplementary Fig. 7 (MSE difference), with *R*^2^ and MSE of DPR.MCMC presented in the corresponding figure legend. Supplementary Table 1 shows the standard deviation of absolute *R*^2^ values across cross variation replicates. Supplementary Fig. 8 also displays trace plots of the log posterior likelihood from DPR.MCMC for all traits, suggesting reasonable convergence of our method.

**Figure 3.**
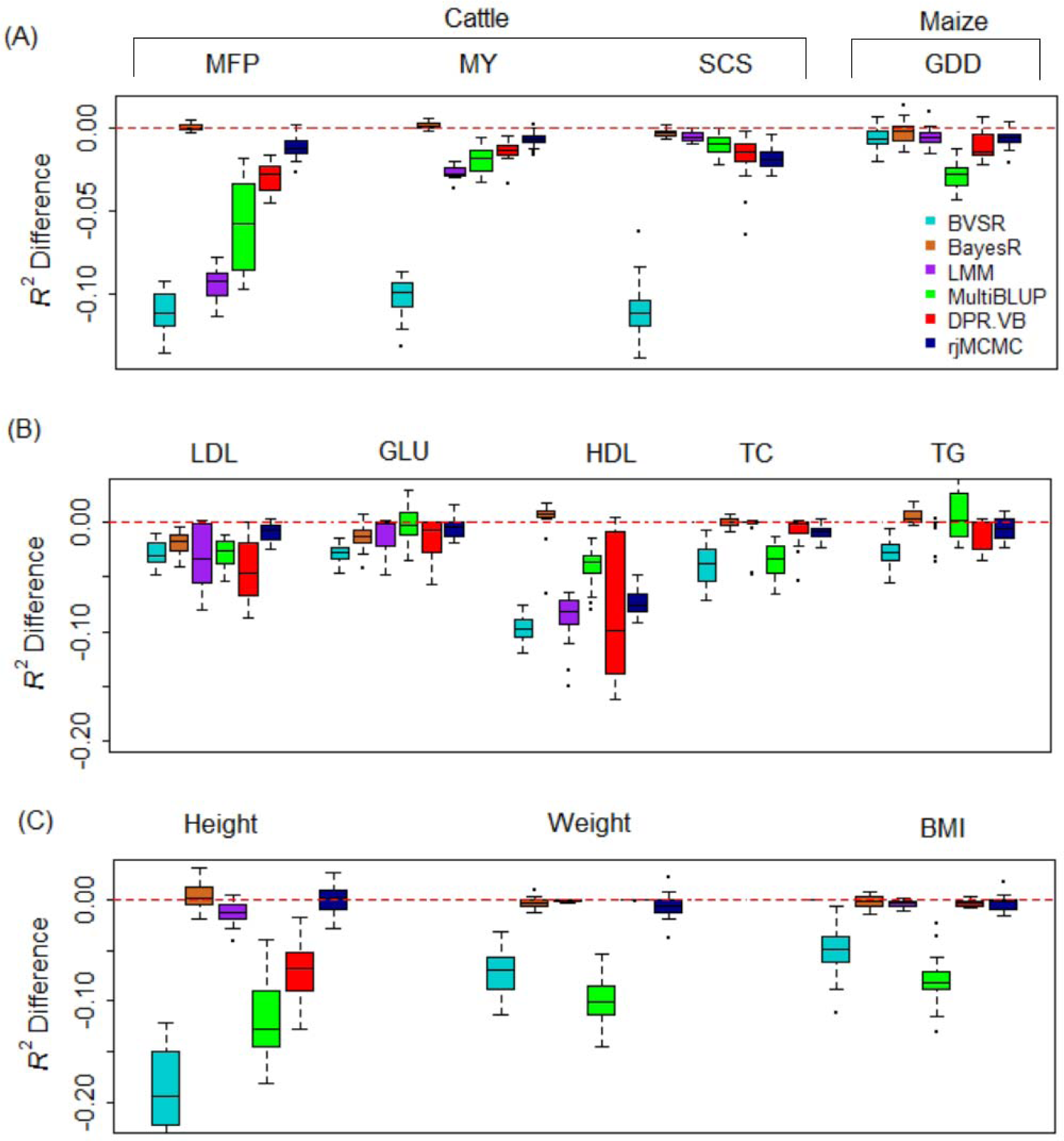
Comparison of prediction performance of several methods with DPR.MCMC for twelve traits from three data sets. Performance is measured by *R*^2^ difference with respect to DPR.MCMC, where a negative value (i.e. values below the red horizontal line) indicates worse performance than DPR.MCMC. Methods for comparison include BVSR (cyan), BayesR (chocolate), LMM (purple), MultiBLUP (green), DPR.VB (red), rjMCMC (black blue) and DPR.MCMC. For each box plot, the bottom and top of the box are the first and third quartiles, while the ends of whiskers represent either the lowest datum within 1.5 interquartile range of the lower quartile or the highest datum within 1.5 interquartile range of the upper quartile. The sample *R*^2^ differences are obtained from 20 replicates of Monte Carlo cross validation for each trait. For DPR.MCMC, the mean predictive *R*^2^ in the test set and the standard deviation across replicates are 0.751 (0.011) for MFP, 0.624 (0.012) for MY, 0.551 (0.017) for SCS and 0.828 (0.012) for GDD, 0.081 (0.033) for LDL, 0.047 (0.017) for GLU, 0.153 (0.044) for HDL, 0.050 (0.020) for TC, 0.044 (0.015) for TG, 0.478 (0.031) for height, 0.169 (0.038) for weight and 0. 137 (0.037) for BMI. The SNP heritability estimates are 0.912 (0.007) for MFP, 0.810 (0.012) for MY, 0.801 (0.012) for SCS, 0.880 (0.013) for GDD, 0.397 (0.024) for LDL, 0.357 (0.036) for GLU, 0.418 (0.024) for HDL, 0.402 (0.036) for TC, 0.334 (0.034) for TG, 0.905 (0.013) for Height, 0.548 (0.022) for Weight and 0.483 (0.023) for BMI.

Overall, consistent with simulations, DPR.MCMC shows robust performance across all traits and is ranked either as the best or the second best method. In the cattle data (Fig. 3A), for MFP and MY, both DPR.MCMC and BayesR perform the best. For SCS, DPR.MCMC performs the best, followed by BayesR. rjMCMC performs well for MFP and MY but poorly for SCS; while LMM and MultiBLUP do not perform well for MFP and MY in the cattle data but their performance improves for SCS. The relative performance of BayesR vs LMM and MultiBLUP in the cattle data is consistent with the distinct genetic architectures that underlie the three complex traits^17,46^: while MFP and MY are affected by a few large or moderate effect SNPs together with many small effect SNPs, SCS is a highly polygenic trait influenced by many SNPs with small effects. BVSR performs poorly for these three traits in the cattle data. In the maize data (Fig. 3A), DPR.MCMC performs the best, followed by BayesR, suggesting that GDD is influenced by a few SNPs with large effects^15^. In the Framingham heart study data (Fig. 3B and Fig. 3C), DPR.MCMC performs the best or among the best for LDL, GLU, TC, Weight and BMI. Its performance is comparable to BayesR and rjMCMC for Height, and follows right behind BayesR for HDL and TG. Besides DPR and BayesR, rjMCMC also performs well in FHS and is often ranked as the third best method (e.g. for LDL, GLU, Height and Weight). In contrast to the relatively robust performance of DPR.MCMC, however, all other methods can perform poorly for certain phenotypes. In Fig. 3, for example, BayesR is the second worst method for predicting GLU; LMM is the second worst method for predicting LDL; MultiBLUP is the worst method for predicting Weight and BMI; DPR.VB performs among the worst for LDL and HDL; rjMCMC performs poorly for HDL; while BVSR performs the worst for almost all traits except for LDL, Height and Weight. The poor performance of BVSR presumably stems from its relatively simple and sparse assumption on the effect sizes.

Because the FHS is a family based study, we use this data to further examine the influence of individual relatedness on prediction performance. To do so, we divided the FHS data into two sub data sets (D1 and D2) with equal sample size but different levels of relatedness (details in Methods): individuals in D1 are more closely related to each other than those in D2. We then compared method performance by performing cross validation in each of the two data sets separately. While the difference between methods becomes smaller due to smaller sample size in the two sub data sets, the relative performance of most methods for the eight traits are largely unchanged in these two sub data sets as compared to that in the complete data (Fig. 9 vs Figs 3B and 3C). For example, DPR.MCMC is ranked as the best method or among the best methods for six traits in D1 and for four traits in D2. BayesR performs similarly and is ranked as the best or among the best for four traits in D1 and for five traits in D2. LMM ranks right after DPR.MCMC and BayesR, while the other methods do not perform well here. In addition, all methods generally perform better in D1 than in D2 (Supplementary Fig. 10), suggesting that relatedness improves prediction performance –– a phenomenon that has been well recognized by previous studies^9,23,47-49^. Besides cross validation within each data set separately, we also performed cross validation between D1 and D2 by predicting traits in one data with parameters inferred from another. The results are again largely consistent with the main results. In particular, DPR.MCMC is ranked as the best or among the best for six traits in D1 to D2 prediction and for eight traits in D2 to D1 prediction. BayesR is ranked as the best or among the best for six traits in D1 to D2 prediction and for eight traits in D2 to D1 prediction. rjMCMC also performs reasonably well and follows right behind DPR.MCMC and BayesR (Supplementary Fig. 11).

### Computational time

Finally, we list the computational time of the seven methods for the twelve traits in Table 3. Note that some differences in computational time among methods may reflect implementation issues, including the language environment in which the methods are implemented, rather than fundamental differences between algorithms. In addition, we only list in the table the computation time in the fitting stage. Computation time spent in the prediction stage by plugging in estimated coefficients in the new data is almost ignorable and is thus not listed. For sampling based methods (BVSR, rjMCMC, DPR.MCMC and BayesR), we measure the computational time based on a fixed number of iterations. However, due to different convergence properties of different algorithms (e.g. BVSR uses a Metropolis-Hastings algorithm, rjMCMC uses a reversible jump MCMC algorithm, while both DPR.MCMC and BayesR use a Gibbs sampling algorithm), a fixed number of iterations in different methods may correspond to different mixing performance. Nevertheless, we can see that DPR.MCMC has a similar computational cost as the other Gibbs based approach (e.g. BayesR), though in the human data both these Gibbs based approaches (DPR.MCMC and BayesR) can be slower than the Metropolis-Hastings approach (BVSR) and the reversible jump MCMC algorithm (rjMCMC) that effectively update only a small subset of significant SNPs in each iteration. In contrast, DPR.VB is orders of magnitude faster than its MCMC counterpart, and is computationally as efficient as the other two non-MCMC based approaches (LMM and MultiBLUP).

**Table 3.**
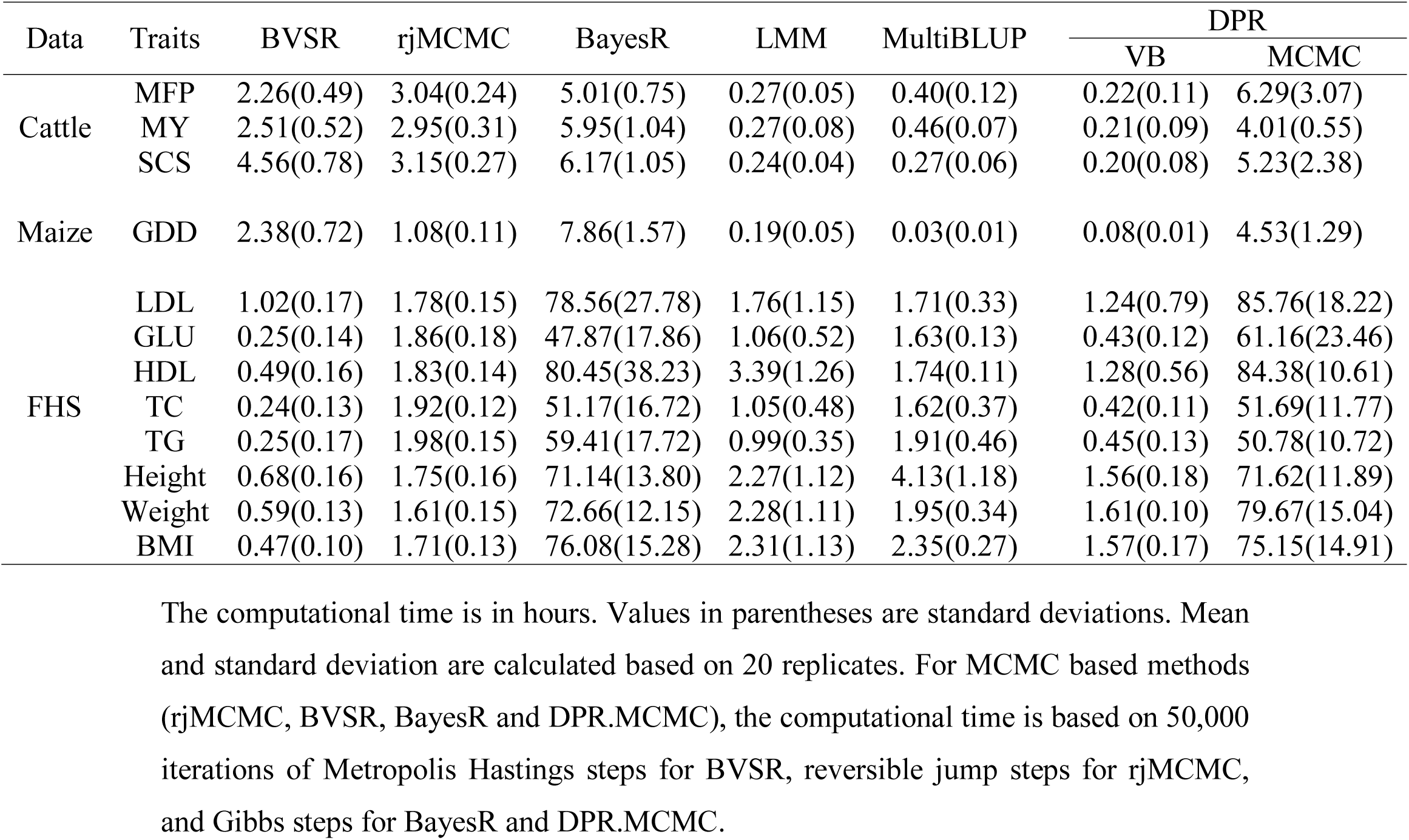
Mean computational time of the seven methods in the model fitting stage for twelve traits across three data sets.

| Data | Traits | BVSR | rjMCMC | BayesR | LMM | MultiBLUP | DPR |  |
| --- | --- | --- | --- | --- | --- | --- | --- | --- |
|  |  |  |  |  |  |  | VB | MCMC |
| Cattle | MFP | 2.26(0.49) | 3.04(0.24) | 5.01(0.75) | 0.27(0.05) | 0.40(0.12) | 0.22(0.11) | 6.29(3.07) |
|  | MY | 2.51(0.52) | 2.95(0.31) | 5.95(1.04) | 0.27(0.08) | 0.46(0.07) | 0.21(0.09) | 4.01(0.55) |
|  | SCS | 4.56(0.78) | 3.15(0.27) | 6.17(1.05) | 0.24(0.04) | 0.27(0.06) | 0.20(0.08) | 5.23(2.38) |
| Maize | GDD | 2.38(0.72) | 1.08(0.11) | 7.86(1.57) | 0.19(0.05) | 0.03(0.01) | 0.08(0.01) | 4.53(1.29) |
| FHS | LDL | 1.02(0.17) | 1.78(0.15) | 78.56(27.78) | 1.76(1.15) | 1.71(0.33) | 1.24(0.79) | 85.76(18.22) |
|  | GLU | 0.25(0.14) | 1.86(0.18) | 47.87(17.86) | 1.06(0.52) | 1.63(0.13) | 0.43(0.12) | 61.16(23.46) |
|  | HDL | 0.49(0.16) | 1.83(0.14) | 80.45(38.23) | 3.39(1.26) | 1.74(0.11) | 1.28(0.56) | 84.38(10.61) |
|  | TC | 0.24(0.13) | 1.92(0.12) | 51.17(16.72) | 1.05(0.48) | 1.62(0.37) | 0.42(0.11) | 51.69(11.77) |
|  | TG | 0.25(0.17) | 1.98(0.15) | 59.41(17.72) | 0.99(0.35) | 1.91(0.46) | 0.45(0.13) | 50.78(10.72) |
|  | Height | 0.68(0.16) | 1.75(0.16) | 71.14(13.80) | 2.27(1.12) | 4.13(1.18) | 1.56(0.18) | 71.62(11.89) |
|  | Weight | 0.59(0.13) | 1.61(0.15) | 72.66(12.15) | 2.28(1.11) | 1.95(0.34) | 1.61(0.10) | 79.67(15.04) |
|  | BMI | 0.47(0.10) | 1.71(0.13) | 76.08(15.28) | 2.31(1.13) | 2.35(0.27) | 1.57(0.17) | 75.15(14.91) |

The computational time is in hours. Values in parentheses are standard deviations. Mean
and standard deviation are calculated based on 20 replicates. For MCMC based methods
(rjMCMC, BVSR, BayesR and DPR.MCMC), the computational time is based on 50,000
iterations of Metropolis Hastings steps for BVSR, reversible jump steps for rjMCMC,
and Gibbs steps for BayesR and DPR.MCMC.

## Discussion

We have presented a novel statistical method, DPR, for genetic prediction of complex traits. DPR uses an infinitely many parameters *a priori* to flexibly model the effect size distribution, and represents the first non-parametric method developed for modeling polygenic traits in genetic association studies. By flexibly modeling the effect size distribution, DPR is capable of adapting to the polygenic architecture underlying many complex traits and enjoys robust performance across a range of phenotypes. With simulations and applications to four real data sets, we have illustrated the benefits of DPR.

We have focused on one application of DPR –– genetic prediction of phenotypes. Like some other polygenic methods^34,35,50^, DPR can also be applied to many other polygenic applications. For example, DPR can be used to estimate the proportion of variance in phenotypes explained by all SNPs, a quantity that is commonly referred to as SNP heritability^28,34^. Because DPR assumes a flexible effect size distribution that is adaptive to the genetic architecture underlying a given trait, it has the potential to provide accurate estimation of SNP heritability. As another example, DPR can be applied to association mapping. There, we can view the normal component with the smallest variance as the polygenic background, and we can estimate the probability of a SNP being in any normal components other than the smallest one as the posterior inclusion probability (PIP). PIP computed in this way measures SNP marginal association strength in the presence of polygenic effects, and may represent a more powerful association indicator than standard single SNP association test statistics^33,50^. An important feature of using PIP in the context of Bayesian models is that PIP quantifies the uncertainty of association strength^33,50^, which is a desirable feature that is not easily achieved by penalized frequentist counterparts^51^.

Here, we have restricted ourselves to applying DPR to continuous phenotypes. For case control studies, we could follow previous approaches of treating binary phenotypes as continuous traits and apply DPR directly^34,35,41^. However, it would be desirable to extend DPR to accommodate case control data or other discrete data types in a principled way, by, for example, extending DPR into the generalized linear model framework. In particular, we could use a probit or a logistic link to extend DPR to directly model case control data. We could use a Poisson or an over-dispersed Poisson distribution to extend DPR to model count data. Extending DPR to various discrete data types would likely lead to wider applications of DPR beyond GWASs. For instance, by modeling count data, DPR could be used to perform differential expression analysis^52^ or expression QTL mapping in RNA sequencing studies^53,54^. Similarly, by modeling proportional data, DPR could be used to perform differential methylation analysis or methyl-QTL mapping in bisulfite sequencing studies^55^. Extending DPR to modeling discrete data types using the generalized linear model framework is thus an important avenue for future research.

In the present study, while we used unrelated individuals for GEUVADIS gene expression prediction and PrediXcan tests, we used related individuals for the other two real data applications. Related individuals not only share similar genetic background but also are likely influenced by a common set of environmental factors^47,56^. In addition, untyped causal SNPs in related individuals can be more easily tagged by neighboring typed SNPs than that in unrelated individuals, thanks to the relatively high linkage disequilibrium (LD) in related data. Because both the shared environmental factors and easy tagging of causal SNPs can facilitate prediction, cross validation using related individuals often results in better prediction performance than using unrelated individuals^9,23,47-49^. However, we caution that the prediction accuracy measured in the test data obtained with cross-validation in related individuals are likely inflated if our ultimate goal is to perform prediction in unrelated individuals instead of related ones. In addition, the predictive model inferred from related individuals may not generalize well to unrelated individuals who are not necessarily influenced by the same set of environmental factors and who do not share the same LD pattern near the causal SNPs. We have attempted to tease apart the influence of relatedness on prediction performance by splitting the FHS data into two parts with different levels of relatedness. Our results indeed show that, while the relative performance of various methods remains largely the same, the absolute performance of all methods do increase with individual relatedness. Additionally, while our method performs well relative to the other methods, we caution that DPR’s prediction accuracy is still unlikely of practical use in human clinical setting. Studies on unrelated individuals or studies using a fully independent validation data are likely required to establish the practical utility of prediction methods, which often have unsatisfactory performance there^9,47,57^. Despite the practical importance of using completely independent or cross-population studies for prediction performance validation, however, we also want to point out its potential caveat: using completely independent data for cross-validation may fail to correctly characterize the relative performance of different methods. In particular, a good method that properly captures the signal in the training data may suffer in the validation data due to different LD patterns between the two data sets. Similarly, a poor method that fails to capture the signal in the training data may perform well in the validation data where such signal is no longer relevant. Therefore, using training and validating data that are both representative of the study population is important to not only ensure a proper comparison among methods but also to ensure the clinical relevance and wide applicability of the prediction methods. Exploring the use of such data is an important direction for future research.

DPR is not without its limitations. Perhaps the biggest limitation is its computational cost. Like any other MCMC based approaches^34,35,58^, our Gibbs algorithm for fitting DPR is computationally slow and can only be applied to moderate-sized GWAS studies. To make DPR widely applicable, we have explored the use of variational Bayesian approximation for fitting DPR. Variational Bayesian approximation obtains an approximate posterior distribution through optimization^59^ and represents a much faster alternative to MCMC sampling. Indeed, DPR.VB is orders of magnitude faster than DPR.MCMC. However, despite its faster computational speed, the VB algorithm is less accurate than MCMC when SNP heritability is large, sometimes by quite a large margin (e.g. PVE=0.80 in simulations). The loss of accuracy in VB is not unexpected because our VB assumes that the posterior distributions of the SNP effect sizes are independent from each other. Posterior independence is an unrealistic assumption given that SNP genotypes are correlated through LD. Therefore, it is important to explore alternative VB algorithms to incorporate the posterior correlation among effect sizes, by, for example, adapting algorithms developed elsewhere^60,61^. It would be ideal if we could develop algorithms that can achieve a high predictive performance as DPR.MCMC but incurs a small computational cost as DPR.VB. Certainly, besides developing alternative algorithms to MCMC, there is still room for improvement on our MCMC algorithm. For example, we could use all individuals to compute some quantities while use only a subset of individuals to compute other quantities, as in our previous MQS method^62^, in order to reduce the computational burden while maintaining the accuracy of the algorithm. In any case, developing efficient and accurate algorithms likely represents a key step to adapt existing polygenic methods to association studies that are orders of magnitude larger.

## Methods

### Overview of DPR

We provide a brief overview of DPR here. Detailed methods and algorithms are provided in the Supplementary Note. To model the relationship between phenotypes and genotypes, we consider the following multiple regression model
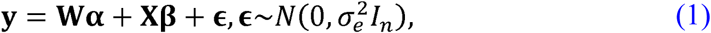

where **y** is an *n*-vector of phenotypes measured on *n* individuals; **W** is an *n* by *c* matrix of covariates including a column of 1 s for the intercept term; **α** is a *c*-vector of coefficients; **X** is an *n* by *p* matrix of genotypes; **β** is the corresponding *p*-vector of effect sizes; ε is an *n*-vector of residual errors where each element is assumed to be independently and identically distributed from a normal distribution with variance 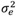.

Like many previous methods^9,19,28,34,41^, we assume that the effect size of *i*th SNP, *β*_*i*_, follows a normal distribution with variance *σ*^2^, i. e. *β_i_*~*N*(0, *σ*^2^). Unlike previous methods, however, we specify a non-parameter prior on the hyper-parameter *σ*^2^ to induce a non-parametric prior on *β*_*i*_. To motivate our prior choice for *σ*^2^, it helps to provide a brief review of the previous polygenic prediction methods. Among the many polygenic prediction methods developed recently, a surprisingly large number of them assume *a priori* that the effect sizes follow a particular class of prior distribution – the scale mixture of normal distributions^63^. Specifically, these methods assume that each effect size *β_i_* follows a normal distribution *β_i_*~*N*(0, *σ*^2^), with the variance parameter (i.e. the scale parameter) *σ*^2^ following another distribution *p*(*σ*^2^). The prior distribution on *σ*^2^, *p*(*σ*^2^), thus differentiates many different predictive methods. For example, LMM assumes a flat prior *p*(*σ*^2^)that is proportional to a constant^9,28^. The Bayes alphabetic methods assume that follows an inverse gamma distribution to induce a t-prior on *β_i_*^10,18,64^. The Bayesian lasso assumes *σ*^2^ that follows a Rayleigh distribution to induce a double exponential distribution (a.k.a. Laplace distribution) on *β_i_*^30,58^. NEG assumes an exponential gamma distribution on *σ*^2^ to induce an NEG prior on *β_i_*^32^. BVSR and BayesCπ assume a mixture of a point mass at zero with another flat prior to induce a point-normal distribution on *β_i_*^29,33^. BSLMM assumes a mixture of two point masses to induce a normal mixture distribution on *β_i_*^34^. While BayesR assumes a three point masses together with another point mass at zero on to also induce a normal mixture distribution on *β_i_*^35^.

The scale mixture of normal distributions is flexible because different distributions on the scale parameter *σ*^2^ can be used to induce many smooth unimodal distributions on *β_i_*. However, existing predictive methods explicitly make a parametric prior assumption on *σ*^2^, which necessarily relies on a limited number of parameters to characterize the distribution on *σ*^2^. Consequently, the induced effect size distribution on *β_i_* from a parametric prior on *σ*^2^ can be restrictive and may sometimes fail to resemble closely the unknown truth effect size distribution underlying complex traits. Motivated by the potential drawback of parametric priors on *σ*^2^, we instead develop a non-parametric prior distribution on *σ*^2^ to induce a more flexible distribution on *β_i_*. Because a non-parametric distribution is characterized by effectively infinitely many parameters, our induced effect size distribution on *β_i_* has the potential to resemble a wide range of genetic architectures and achieve robust predictive performance across a variety of traits.

Technically, we assume *σ*^2^ follows a Dirichlet process (DP) prior^37-40^
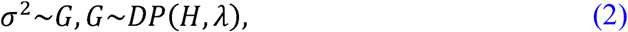

where *H* is the base distribution, and λ is the concentration parameter that describes how the distribution on *σ*^2^, *G*, deviates from the base distribution. Here, we use an inverse gamma distribution as the base distribution and set the two parameters in the inverse gamma distribution to small values to keep the prior relatively uninformative. We treat the concentration parameter *λ* as an unknown hyper-parameter and intend to infer it from the data at hand. Because we use the Dirichlet process as a prior for the latent variance parameter *σ*^2^ we refer to our regression model based on equations (1)-(2) as the latent Dirichlet Process Regression, or DPR. The induced marginal distribution on *β_i_* (after integrating out *σ*^2^) is also non-parametric and can robustly resemble a large classes of unimodal distributions. Indeed, the distribution on *β_i_* can be used to adaptively and accurately approximate a t-distribution, a point-t mixture distribution, a mixture of step functions, as well as the marginal effect sizes estimated from a real data set; whereas a normal distribution cannot (Fig. 1). Therefore, our prior distribution on the effect size can adaptively approximate a wide range of possible effect size distributions underlying complex traits. Since accurate modeling of the effect size distribution is a key to achieve accurate prediction performance^24,34,36^, we expect our non-parametric model to perform robustly well across a range of polygenic architectures.

It is important to point out that our modeling assumption on the effect sizes *β_i_* is also mathematically equivalent to a Dirichlet process normal mixture, which is a mixture of normal distributions with infinitely many normal components. Specifically, using the stick-breaking constructive representation of the Dirichlet process^59^, we can re-write our modeling assumption on *β_i_* in an equivalent form as
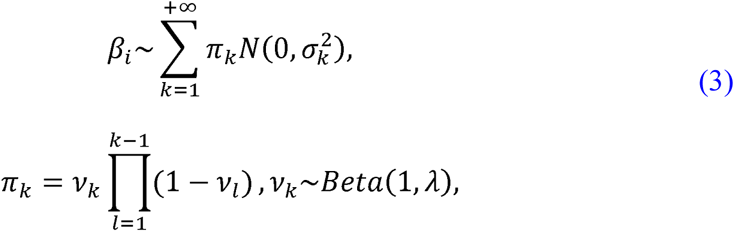

where *λ* is the same concentration parameter as in equation (2), and determines the number of normal components in the model and subsequently the model complexity^59^. Each 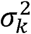 in the above equation follows the base distribution *H*. From the normal mixture equivalence aspect, our method effectively generalizes many previous methods^18,34,35^ that use a fixed, often small, number of normal components to using infinitely many normal components *a priori*. Although the prior number of normal components in our model is infinite, the posterior number of components for any given data set will be finite, and can be automatically inferred based on the data at hand. Therefore, our model has the potential to adjust the model complexity according to the data complexity, and has the potential to adapt to a wide range of polygenic architectures.

To fit our model, we develop two complementary algorithms: one is based on the MCMC algorithm, and the other is based on the variational Bayesian (VB) approximation. The MCMC sampling algorithm, which we refer to as DPR.MCMC, is accurate but computationally slow. The variational Bayesian algorithm, which we refer to as DPR.VB, is computationally fast, but, as we will show in the results, is often less accurate. The two algorithms provide users the choice of speed vs accuracy. The details of the two algorithms are provided in Supplementary Note.

### Simulations

we used genotypes from an existing cattle GWAS dataset^17^ with 5,024 individuals and 42,551 SNPs and simulated phenotypes. To cover a range of possible genetic architectures, we consider *eight* simulation settings from four different simulation scenarios to cover a range of possible genetic architectures:

1. Scenario I satisfies the DPR modeling assumption, where all SNPs are causal and SNPs in different effect-size groups have different effects. Specifically, we randomly selected 10 group-one SNPs, 100 group-two SNPs, 1,000 group-three SNPs, and set the remaining SNPs as group-four SNPs. We simulated SNP effect sizes all from a standard normal distribution but scaled their effects in each group separately so that the proportion of genetic variance explained by the four groups are 0.05, 0.15, 0.20, and 0.60, respectively. We set the total proportion of phenotypic variance (PVE; i.e. SNP heritability) to be either 0.2, 0.5 or 0.8, representing low, moderate, and high heritability, respectively. This simulation scenario consists of one simulation setting for each PVE.
2. Scenario II satisfies the BayesR modeling assumption, where a small proportion of SNPs are causal. These causal SNPs come from three effect-size groups. The simulations were similar to scenario I with the only exception that the group-four SNPs have zero effects. Here, the proportion of PVE by the three groups are 0.1, 0.2, and 0.7, respectively. Again, we set the total PVE to be either 0.2, 0.5 or 0.8. This simulation scenario consists of one simulation setting for each PVE.
3. Scenario III is similar to Scenario I except that SNPs come from two effect-size groups, thus representing a simpler scenario than I. In particular, we selected either c=10, 100 or 1,000 SNPs as group-one SNPs and set the remaining SNPs as group-two SNPs. We simulated their effect sizes from a standard normal distribution and scaled their effects in each group separately so that the proportion of PVE by the two groups are 0.2 and 0.8, respectively. Again, we set the total PVE to be either 0.2, 0.5 or 0.8. This simulation scenario consists of three simulation settings for each PVE (c=10, 100 or 1,000).
4. Scenario IV is related to the assumption made in LMM and MultiBLUP. Here, all SNPs have non-zero effects and their effect sizes come from either a normal distribution, a t-distribution with four degrees of freedom, or a Laplace distribution. We scaled their effect sizes further so that the total PVE equals 0.2, 0.5, or 0.8. This simulation scenario consists of three simulation settings for each PVE (normal, t, or Laplace).

In each setting, we performed 20 simulation replicates. In each replicate, we randomly split the data into a training data with 80% individuals and a test data with the remaining 20% individuals. We then fitted different methods on the training data and evaluated their prediction performance on the test data (i.e. Monte Carlo cross validation).

### GEUVADIS data

The GEUVADIS data^42^ contains gene expression measurements for 465 individuals from five different populations: CEPH (CEU), Finns (FIN), British (GBR), Toscani (TSI) and Yoruba (YRI). Following previous studies^65^, we focused only on protein coding genes and lincRNAs that are annotated from GENCODE^66^ (release 12). We removed lowly expressed genes that have zero counts in at least half of the individuals and obtained a final set of 15,810 genes. Afterwards, following previous studies^67^, we performed PEER normalization to remove confounding effects and unwanted variations. In order to remove potential population stratification, we quantile normalized the gene expression measurements across individuals in each population to a standard normal distribution, and then quantile normalized the gene expression measurements to a standard normal distribution across individuals from all five populations. In addition to the gene expression data, all individuals in GEUVADIS also have their genotypes sequenced in the 1000 Genomes Project. Among the sequenced genotypes, we filtered out SNPs that have a Hardy-Weinberg equilibrium (HWE) p-value < 10^-4^, a genotype call rate < 95%, or an MAF < 0.01. We retained a total of 7,072,917 SNPs for analysis. We intersected these SNPs with imputed SNPs from WTCCC data^4^ (see below; for the purpose of performing gene set tests) and kept a final set of 2,793,818 overlapping SNPs for analysis. Then, for each gene in turn, we obtained its cis-SNPs that are located within either 100 kb upstream of the transcription start site (TSS) or 100 kb downstream of the transcription end site (TES), resulting in an average of 175 cis-SNPs per gene.

### WTCCC data

The Wellcome Trust Case Control Consortium^4^ (WTCCC) 1 data consists of about 14,000 cases from seven common diseases and 2,938 shared controls. The cases include 1,963 individuals with type 1 diabetes (T1D), 1,748 individuals with Crohn’s disease (CD), 1,860 individuals with rheumatoid arthritis (RA), 1,868 individuals with bipolar disorder (BD), 1,924 individuals with type 2 diabetes (T2D), 1,926 individuals with coronary artery disease (CAD), and 1,952 individuals with hypertension (HT). We obtained quality controlled genotypes from WTCCC and imputed missing genotypes using BIMBAM^68^. We obtained a total of 458,868 SNPs shared across all individuals. We then further imputed SNPs using the 1000 Genomes as the reference panel with SHAPEIT and IMPUTE2^69^. We filtered out SNPs that have a HWE p-value < 10^-4^, a genotype call rate < 95%, or an MAF < 0.01 to obtain a total of 2,793,818 imputed SNPs. For PrediXcan analysis^12^, as in the GEUVADIS data (see above), we focused on the same 15,810 genes. As in^12^, we further restricted our association analysis on a set of 4,343 genes that have a predictive *R*^2^ above 0.01 by all predictive methods.

### Cattle data

The cattle data^17^ consists of 5,024 samples and 42,551 SNPs after removing SNPs that have a HWE p-value < 10^-4^, a genotype call rate < 95%, or an MAF < 0.01. For the remaining SNPs, we imputed missing genotypes with the estimated mean genotype of that SNP. We analyzed three traits: milk fat percentage (MFP), milk yield (MY), and somatic cell score (SCS). All phenotypes were quantile normalized to a standard normal distribution before analysis.

### Maize data

The maize data^15^ consists of 2,267 inbred accessions and 98,385 SNPs after removing SNPs that have a HWE p-value < 10^-4^, a genotype call rate < 95%, or an MAF < 0.01. For the remaining SNPs, we imputed missing genotypes with the estimated mean genotype of that SNP. We used the growing degree days (GDD) to silking as the phenotype in genomic selection. GDD was calculated using climate data from weather stations located near the farms^15^, and was quantile normalized to a standard normal distribution before analysis.

### Framingham heart study data

The Framingham heart study (FHS) data contains genotype data on 6,950 individuals and 394,174 SNPs. We filtered out SNPs that have a HWE p-value < 10^-4^, a genotype call rate < 95%, or an MAF < 0.01 to obtain a final set of 387,741 SNPs. For these SNPs, we imputed missing genotypes with the estimated mean genotype of that SNP. We performed analysis on eight traits: five commonly used plasma traits that include low-density lipoprotein (LDL) cholesterol, glucose (GLU), high-density lipoprotein (HDL) cholesterol, total cholesterol (TC), and triglycerides (TG); and three anthropometric traits that include height, weight, and body mass index (BMI). Each trait was quantile normalized to a standard normal distribution before analysis. Note that the FHS data is a family-based study where individuals are genetically related. To tease apart the influence of individual relatedness on prediction performance among methods, we also divided the samples in FHS into two separate data sets with different levels of relatedness. Specifically, we first used genotypes to compute the genome-wide genetic relatedness matrix (GRM). We then ordered individual pairs based on their genetic relatedness values. From top to bottom of the ordered individual pair list, we selected individuals from individual pairs with high levels of relatedness into a data set D1, and continued this process until the sample size of D1 was half of the full data. We then kept the remaining individuals from individual pairs with low levels of relatedness into a data set D2. The relatedness threshold for separating individual pairs between the two data sets was 0.151. Nevertheless, the majority pairs in D1 and D2 have genetic relatedness values close to zero: 99.6% of pairs in D1 and 99.9% of pairs in D2 have a genetic relatedness value between +/-0.01. As another way of measuring relatedness, we also computed the effective number of chromosome segments (*M*_e_)^49^ in the two data. *M*_e_ is a crucial parameter that measures the effective number of independent SNPs and is also closely related to the effective number of independent individuals: *M*_e_ is small in a data with related individuals and is large in a data with unrelated individuals. A small value of *M*_e_ often correlates to high prediction accuracy^48,49,70^. With 20 cross-validation replicates, we estimated *M*_e_ in D1 and D2 sub data to be 34541.39 (sd=140.87) and 81786.01 (sd=651.52), respectively.

### Other methods

We compared the performance of DPR.MCMC and DPR.VB mainly with five existing methods: (1) LMM^28^ as implemented in the GEMMA software (version 0.95alpha); (2) BVSR^29^ as implemented in the GEMMA software (version 0.95alpha); (3) MultiBLUP^41^ as implemented in the LDAK software (version 4.9); (4) BayesR^35^ as implemented in the bayesR software; (5) rjMCMC^20^ as implemented in the gwas_rjmc 1.163 software. We used default settings to fit all these methods. For rjMCMC, because it requires us to provide a variance component parameter, we used LMM to estimate the variance component first in all analyses. In addition, rjMCMC does not output parameter estimates. Therefore, for the PrediXcan analysis, we first merged the GEUVADIS and WTCCC files for every gene, labeled WTCCC individuals as having missing phenotypes, and then ran rjMCMC on these files to obtained predicted gene expression values using the WTCCC genotype data. The same strategy was also applied to perform cross-validation prediction between D1 and D2 sub data sets in FHS. For gene expression prediction and PrediXcan analysis, following the original PrediXcan paper^12^, we also used Elastic Net (ENET)^44^, which is implemented in the R package glmnet (version 1.9-5). For ENET, following^12^, we set one penalty parameter (i.e. *α*) to be 0.5 and selected the other one using 100-fold cross validation in the training data.

### Data availability

No data were generated in the present study. The GEUVADIS gene expression data is publicly available at http://www.geuvadis.org. The genotype data from the 1000 genomes project is publicly available at http://www.internationalgenome.org. The WTCCC genotype and phenotype data is publicly available at https://www.wtccc.org.uk. The genotype and phenotype data from the cattle and maize studies are available from the authors upon reasonable request and with permission of Prof. Xiaolei Liu at the HuaZhong Agriculture University. The Framingham heart study genotype and phenotype data is available in dbGaP (https://www.ncbi.nlm.nih.gov/gap) with accession number phs000007.

### Software availability

Our method is implemented in the DPR software, freely available at http://www.xzlab.org/software.html.

## Acknowledgments

This research is supported by National Institutes of Health grant R01HG009124. XZ is also supported by NIH Grants R01HL117626 (PI Abecasis), R21ES024834 (PI Pierce), R01HL133221 (PI Smith), and a grant from the Foundation for the National Institutes of Health through the Accelerating Medicines Partnership (BOEH15AMP, co-PIs Boehnke and Abecasis). We thank Xiaolei Liu from the HuaZhong Agriculture University for providing us with the cattle and maize data. We thank Dr. Shiquan Sun for help with the development of DPR. We thank Dr. Doug Speed for help with LDAK and Dr. Sang Hong Lee for help with gwas_rjmc1.163 software. This study also makes use of data generated by the Wellcome Trust Case Control Consortium (WTCCC). A full list of the investigators who contributed to the generation of the data is available from http://www.wtccc.org.uk. Funding for the WTCCC project was provided by the Wellcome Trust under award 076113 and 085475. This research was conducted in part using data and resources from the Framingham Heart Study of the NHLBI and Boston University School of Medicine, which was partially supported by the NHLBI Framingham Heart Study (Contract No. N01-HC-25195) and its contract with Affymetrix, Inc for genotyping services (Contract No. N02-HL-6-4278). We thank all participants and staff from the Framingham Heart Study. We thank the three anonymous reviewers for their constructive comments which greatly improved the quality of the paper.

## Author contributions

X.Z. and P.Z. conceived and designed the experiments, developed the algorithm and implemented the software used in analysis. P.Z. performed the experiments, analyzed the data. X.Z. and P.Z. wrote the paper.

## Additional information

### Supplementary Information

**Supplementary Figure 1. Comparison of prediction performance of several methods with DPR.MCMC in simulations when PVE=0.2.** Performance is measured by *R*^2^ difference with respect to DPR.MCMC, where a negative value (i.e. values below the red horizontal line) indicates worse performance than DPR.MCMC. The sample *R*^2^ differences are obtained from 20 replicates in each scenario. Methods for comparison include BVSR (cyan), BayesR (chocolate), LMM (purple), MultiBLUP (green), DPR.VB (red), rjMCMC (black blue) and DPR.MCMC. Simulation scenarios include: (A) Scenario I, which satisfies the DPR modeling assumption; (B) Scenario II, which satisfies the BayesR modeling assumption; (C) Scenario III, where the number of SNPs in the large effect group is 10, 100, or 1,000; and (D) Scenario IV, where the effect sizes are generated from either a normal distribution, a t-distribution or a Laplace distribution. For each box plot, the bottom and top of the box are the first and third quartiles, while the ends of whiskers represent either the lowest datum within 1.5 interquartile range of the lower quartile or the highest datum within 1.5 interquartile range of the upper quartile. For DPR.MCMC, the mean predictive *R*^2^ in the test set and the standard deviation for the eight settings are respectively 0.074 (0.020), 0.081 (0.016), 0.076 (0.018), 0.072 (0.019), 0.064 (0.016), 0.083 (0.023), 0.077 (0.016) and 0.077 (0.017).

**Supplementary Figure 2. Comparison of prediction performance of several methods with DPR.MCMC in simulations when PVE=0.8.** Performance is measured by *R*^2^ difference with respect to DPR.MCMC, where a negative value (i.e. values below the red horizontal line) indicates worse performance than DPR.MCMC. The sample *R*^2^ differences are obtained from 20 replicates in each scenario. Methods for comparison include BVSR (cyan), BayesR (chocolate), LMM (purple), MultiBLUP (green), DPR.VB (red), rjMCMC (black blue) and DPR.MCMC. Simulation scenarios include: (A) Scenario I, which satisfies the DPR modeling assumption; (B) Scenario II, which satisfies the BayesR modeling assumption; (C) Scenario III, where the number of SNPs in the large effect group is 10, 100, or 1,000; and (D) Scenario IV, where the effect sizes are generated from either a normal distribution, a t-distribution or a Laplace distribution. For each box plot, the bottom and top of the box are the first and third quartiles, while the ends of whiskers represent either the lowest datum within 1.5 interquartile range of the lower quartile or the highest datum within 1.5 interquartile range of the upper quartile. For DPR.MCMC, the mean predictive *R*^2^ in the test set and the standard deviation for the eight settings are respectively 0.554 (0.028), 0.622 (0.022), 0.569 (0.023), 0.548 (0.027), 0.537 (0.030), 0.543 (0.028), 0.546 (0.027) and 0.539 (0.022).

**Supplementary Figure 3. Comparison of prediction performance of several methods with DPR.MCMC in simulations when PVE=0.2.** Performance is measured by MSE difference with respect to DPR.MCMC, where a positive value (i.e. values above the red horizontal line) indicates worse performance than DPR.MCMC. The sample MSE differences are obtained from 20 replicates in each scenario. Methods for comparison include BVSR (cyan), BayesR (chocolate), LMM (purple), MultiBLUP (green), DPR.VB (red), rjMCMC (black blue) and DPR.MCMC. Simulation scenarios include: (A) Scenario I, which satisfies the DPR modeling assumption; (B) Scenario II, which satisfies the BayesR modeling assumption; (C) Scenario III, where the number of SNPs in the large effect group is 10, 100, or 1,000; and (D) Scenario IV, where the effect sizes are generated from either a normal distribution, a t-distribution or a Laplace distribution. For each box plot, the bottom and top of the box are the first and third quartiles, while the ends of whiskers represent either the lowest datum within 1.5 interquartile range of the lower quartile or the highest datum within 1.5 interquartile range of the upper quartile. For DPR.MCMC, the mean predictive MSE in the test set and the standard deviation for the eight settings are respectively 0.919 (0.044), 0.910 (0.038), 0.929 (0.036), 0.944 (0.053), 0.923 (0.038), 0.925 (0.033), 0.924 (0.037) and 0.918 (0.037).

**Supplementary Figure 4. Comparison of prediction performance of several methods with DPR.MCMC in simulations when PVE=0.5.** Performance is measured by MSE difference with respect to DPR.MCMC, where a positive value (i.e. values above the red horizontal line) indicates worse performance than DPR.MCMC. The sample MSE differences are obtained from 20 replicates in each scenario. Methods for comparison include BVSR (cyan), BayesR (chocolate), LMM (purple), MultiBLUP (green), DPR.VB (red), rjMCMC (black blue) and DPR.MCMC. Simulation scenarios include: (A) Scenario I, which satisfies the DPR modeling assumption; (B) Scenario II, which satisfies the BayesR modeling assumption; (C) Scenario III, where the number of SNPs in the large effect group is 10, 100, or 1,000; and (D) Scenario IV, where the effect sizes are generated from either a normal distribution, a t-distribution or a Laplace distribution. For each box plot, the bottom and top of the box are the first and third quartiles, while the ends of whiskers represent either the lowest datum within 1.5 interquartile range of the lower quartile or the highest datum within 1.5 interquartile range of the upper quartile. For DPR.MCMC, the mean predictive MSE in the test set and the standard deviation for the eight settings are respectively 0.722 (0.043), 0.701 (0.028), 0.707 (0.034), 0.717 (0.037), 0.727 (0.034), 0.734 (0.040), 0.721 (0.032) and 0.720 (0.028).

**Supplementary Figure 5. Comparison of prediction performance of several methods with DPR.MCMC in simulations when PVE=0.8.** Performance is measured by MSE difference with respect to DPR.MCMC, where a positive value (i.e. values above the red horizontal line) indicates worse performance than DPR.MCMC. The sample MSE differences are obtained from 20 replicates in each scenario. Methods for comparison include BVSR (cyan), BayesR (chocolate), LMM (purple), MultiBLUP (green), DPR.VB (red), rjMCMC (black blue) and DPR.MCMC. Simulation scenarios include: (A) Scenario I, which satisfies the DPR modeling assumption; (B) Scenario II, which satisfies the BayesR modeling assumption; (C) Scenario III, where the number of SNPs in the large effect group is 10, 100, or 1,000; and (D) Scenario IV, where the effect sizes are generated from either a normal distribution, a t-distribution or a Laplace distribution. For each box plot, the bottom and top of the box are the first and third quartiles, while the ends of whiskers represent either the lowest datum within 1.5 interquartile range of the lower quartile or the highest datum within 1.5 interquartile range of the upper quartile. For DPR.MCMC, the mean predictive MSE in the test set and the standard deviation for the eight settings are respectively 0.443 (0.032), 0.379 (0.016), 0.429 (0.024), 0.454 (0.023), 0.464 (0.030), 0.465 (0.027), 0.454 (0.032) and 0.457 (0.022).

**Supplementary Figure 6. Comparison of predictive *R*^2^ from DPR.MCMC with the other six methods for predicting gene expression levels in the GEUVADIS data.** Scatter plots show (A) predictive *R*^2^ in the test data obtained by DPR.MCMC vs that obtained by BVSR for all genes; (B) DPR.MCMC vs ENET; (C) DPR.MCMC vs BayesR; (D) DPR.MCMC vs LMM; (E) DPR.MCMC vs MultiBLUP; (F) DPR.MCMC vs DPR.VB; (G) DPR.MCMC vs rjMCMC. Each panel also lists the number of genes where DPR.MCMC performs better (first number) and the number of genes where DPR.MCMC performs worse (second number).

**Supplementary Figure 7. Comparison of prediction performance of several methods with DPR.MCMC for twelve traits from three data sets.** Performance is measured by MSE difference with respect to DPR.MCMC, where a positive value (i.e. values above the red horizontal line) indicates worse performance than DPR.MCMC. Methods for comparison include BVSR (cyan), BayesR (chocolate), LMM (purple), MultiBLUP (green), DPR.VB (red), rjMCMC (black blue) and DPR.MCMC. The sample MSE differences are obtained from 20 replicates of Monte Carlo cross validation for each trait. For each box plot, the bottom and top of the box are the first and third quartiles, while the ends of whiskers represent either the lowest datum within 1.5 interquartile range of the lower quartile or the highest datum within 1.5 interquartile range of the upper quartile. For DPR.MCMC, the mean predictive MSE in the test set and the standard deviation are 0.246 (0.011) for MFP, 0.371 (0.019) for MY, 0.446 (0.028) for SCS, 0.170 (0.012) for GDD, 0.928 (0.029) for LDL, 0.954 (0.034) for GLU, 0.833 (0.063) for HDL, 0.970 (0.044) for TC, 0.960 (0.035) for TG, 0.519 (0.050) for height, 0.834 (0.065) for weight and 0.868 (0.074) for BMI. The SNP heritability estimates are 0.912 (0.007) for MFP, 0.810 (0.012) for MY, 0.801 (0.012) for SCS, 0.880 (0.013) for GDD, 0.397 (0.024) for LDL, 0.357 (0.036) for GLU, 0.418 (0.024) for HDL, 0.402 (0.036) for TC, 0.334 (0.034) for TG, 0.905 (0.013) for Height, 0.548 (0.022) for Weight and 0.483 (0.023) for BMI.

**Supplementary Figure 8. Trace plots of the log posterior likelihood of DPR.MCMC in real data applications.** For each of the twelve traits in the three GWAS data sets, we plot the log posterior likelihood versus the first 10,000 iterations (i.e. burn-in period) using the first cross-validation data. In each panel, the log posterior likelihood values were centered to have a median value of zero.

**Supplementary Figure 9. Comparison of prediction performance of several methods with DPR.MCMC for eight traits in each of the two sub data sets of FHS.** The two sub data sets D1 and D2 have the same sample size but different levels of relatedness (individuals in D1 are more related to each other than those in D2). (A) The *R*^2^ difference of five plasma traits (LDL, GLU, HDL, TC and TG) with respect to DPR.MCMC in the D1 and D2 sub data of FHS; (B) The *R*^2^ difference of three anthropometric traits (Height, Weight and BMI) with respect to DPR.MCMC in the D1 and D2 sub data of FHS. For each box plot, the bottom and top of the box are the first and third quartiles, while the ends of whiskers represent either the lowest datum within 1.5 interquartile range of the lower quartile or the highest datum within 1.5 interquartile range of the upper quartile. FHS: Framingham heart study.

**Supplementary Figure 10. Prediction performance of various methods are higher in a data with more related individuals (D1) than in a data with less related individuals (D2).** The two data sets D1 and D2 from FHS have the same sample size but different levels of relatedness (individuals in D1 are more related to each other than those in D2). For each trait in the FHS data (x-axis), we first computed the median predictive *R*^2^ across 20 replicates in D1 and D2 separately, and then contrast the difference between the two averaged predictive *R*^2^ values in the two data sets (D1 minus D2; y-axis). Positive averaged predictive *R*^2^ differences suggest that all methods have higher predictive performance in D1 versus D2. FHS: Framingham heart study.

**Supplementary Figure 11. Comparison of prediction performance of several methods with DPR.MCMC using cross-validation between the two sub data sets of FHS.** The two sub data sets D1 and D2 have the same sample size but different levels of relatedness (individuals in D1 are more related to each other than those in D2). (A) Predictive *R*^2^ difference of different methods in D1 using parameters inferred in D2. For DPR.MCMC, the *R*^2^ is 0.024 for LDL, 0.012 for GLU, 0.021 for HDL, 0.022 for TC, 0.016 for TG, 0.131 for Height, 0.061 for Weight and 0.041 for BMI. (B) Predictive *R*^2^ difference of different methods in D2 using parameters inferred in D1; For DPR.MCMC, the *R*^2^ is 0.043 for LDL, 0.009 for GLU, 0.033 for HDL, 0.021 for TC, 0.015 for TG, 0.226 for Height, 0.083 for Weight and 0.058 for BMI. FHS: Framingham heart study.

**Supplementary Table 1. Sampling variation of *R*^2^ measured by standard deviation across Monte Carlo cross validation replicates for various methods in simulations and real data analysis.**

**Supplementary Table 2. Significant genes identified by DPR.MCMC for different diseases in the PrediXcan gene set analysis of WTCCC.**

**Supplementary Note. Model and Algorithm Details for DPR**

## References

1. Fritsche, L. G. et al. A large genome-wide association study of age-related macular degeneration highlights contributions of rare and common variants. Nat. Genet. 48, 134-143 (2016).

2. Mancuso, N. et al. The contribution of rare variation to prostate cancer heritability. Nat. Genet. 48, 30-35 (2016).

3. Fuchsberger, C. et al. The genetic architecture of type 2 diabetes. Nature 536, 41-47 (2016).

4. The Wellcome Trust Case Control Consortium. Genome-wide association study of 14,000 cases of seven common diseases and 3,000 shared controls. Nature 447, 661-678 (2007).

5. Global Lipids Genetics Consortium. Discovery and refinement of loci associated with lipid levels. Nat. Genet. 45, 1274-1283 (2013).

6. Afshari, N. A. et al. Genome-wide association study identifies three novel loci in Fuchs endothelial corneal dystrophy. Nat. Commun. 8, 14898 (2017).

7. Hoffmann, T. J. et al. Genome-wide association study of prostate-specific antigen levels identifies novel loci independent of prostate cancer. Nat. Commun. 8, 14248 (2017).

8. Warren, H. R. et al. Genome-wide association analysis identifies novel blood pressure loci and offers biological insights into cardiovascular risk. Nat. Genet. 49, 403-415 (2017).

9. Makowsky, R. et al. Beyond Missing Heritability: Prediction of Complex Traits. PLoS Genet. 7, e1002051 (2011).

10. Hayes, B. J., Pryce, J., Chamberlain, A. J., Bowman, P. J., & Goddard, M. E. Genetic Architecture of Complex Traits and Accuracy of Genomic Prediction: Coat Colour, Milk-Fat Percentage, and Type in Holstein Cattle as Contrasting Model Traits. PLoS Genet. 6, e1001139 (2010).

11. Chatterjee, N., Shi, J., & Garcia-Closas, M. Developing and evaluating polygenic risk prediction models for stratified disease prevention. Nat. Rev. Genet. 17, 392-406 (2016).

12. Gamazon, E. R. et al. A gene-based association method for mapping traits using reference transcriptome data. Nat. Genet. 47, 1091-1098 (2015).

13. Allen, H. L. et al. Hundreds of variants clustered in genomic loci and biological pathways affect human height. Nature 467, 832-838 (2010).

14. Li, H. et al. Genome-wide association study dissects the genetic architecture of oil biosynthesis in maize kernels. Nat. Genet. 45, 43-50 (2013).

15. Romay, M. C. et al. Comprehensive genotyping of the USA national maize inbred seed bank. Genome Biol. 14, R55 (2013).

16. Fernandes Júnior, G. A. et al. Genomic prediction of breeding values for carcass traits in Nellore cattle. Genet. Sel. Evol. 48, 7 (2016).

17. Zhang, Z. et al. Accuracy of whole-genome prediction using a genetic architecture-enhanced variance-covariance matrix. G3: Genes | Genomes | Genetics 5, 615-627 (2015).

18. Meuwissen, T., Hayes, B., & Goddard, M. Prediction of total genetic value using genome-wide dense marker maps. Genetics 157, 1819-1829 (2001).

19. de los Campos, G., Vazquez, A. I., Fernando, R., Klimentidis, Y. C., & Sorensen, D. Prediction of complex human traits using the genomic best linear unbiased predictor. PLoS Genet. 9, e1003608 (2013).

20. Lee, S. H., van der Werf, J. H. J., Hayes, B. J., Goddard, M. E., & Visscher, P. M. Predicting Unobserved Phenotypes for Complex Traits from Whole-Genome SNP Data. PLoS Genet. 4, e1000231 (2008).

21. Hayes, B., Bowman, P., Chamberlain, A., & Goddard, M. Genomic selection in dairy cattle: Progress and challenges. J. Dairy Sci. 92, 433-443 (2009).

22. Goddard, M. E., & Hayes, B. Genomic selection. J. Anim. Breed. Genet. 124, 323-330 (2007).

23. Meuwissen, T., Hayes, B., & Goddard, M. Accelerating improvement of livestock with genomic selection. Annu. Rev. Anim. Biosci. 1, 221-237 (2013).

24. Chatterjee, N. et al. Projecting the performance of risk prediction based on polygenic analyses of genome-wide association studies. Nat. Genet. 45, 400-405 (2013).

25. Shah, S. et al. Improving Phenotypic Prediction by Combining Genetic and Epigenetic Associations. Am. J. Hum. Genet. 97, 75-85 (2015).

26. Maier, R. et al. Joint analysis of psychiatric disorders increases accuracy of risk prediction for schizophrenia, bipolar disorder, and major depressive disorder. Am. J. Hum. Genet. 96, 283-294 (2015).

27. Weissbrod, O., Geiger, D., & Rosset, S. Multikernel: linear mixed models for complex phenotype prediction. Genome Res. 26, 969-979 (2016).

28. Yang, J. et al. Common SNPs explain a large proportion of the heritability for human height. Nat. Genet. 42, 565-569 (2010).

29. Habier, D., Fernando, R. L., Kizilkaya, K., & Garrick, D. J. Extension of the bayesian alphabet for genomic selection. BMC Bioinformatics 12, 186 (2011).

30. Park, T., & Casella, G. The Bayesian Lasso. J. Am. Stat. Assoc. 103, 681-686 (2008).

31. Yi, N., & Xu, S. Bayesian LASSO for Quantitative Trait Loci Mapping. Genetics 179, 1045-1055 (2008).

32. Hoggart, C. J., Whittaker, J. C., De Iorio, M., & Balding, D. J. Simultaneous analysis of all SNPs in genome-wide and re-sequencing association studies. PLoS Genet. 4, e1000130 (2008).

33. Guan, Y., & Stephens, M. Bayesian variable selection regression for genome-wide association studies and other large-scale problems. Ann. Appl. Stat. 5, 1780-1815 (2011).

34. Zhou, X., Carbonetto, P., & Stephens, M. Polygenic modeling with Bayesian sparse linear mixed models. PLoS Genet. 9, e1003264 (2013).

35. Moser, G. et al. Simultaneous Discovery, Estimation and Prediction Analysis of Complex Traits Using a Bayesian Mixture Model. PLoS Genet. 11, e1004969 (2015).

36. Goddard, M. Genomic selection: prediction of accuracy and maximisation of long term response. Genetica 136, 245-257 (2009).

37. Ghahramani, Z. Bayesian non-parametrics and the probabilistic approach to modelling. Philos. T. R. Soc. A 371, 20110553 (2013).

38. Müller, P., & Mitra, R. Bayesian Nonparametric Inference—Why and How. Bayesian. Anal. 8, 269-302 (2013).

39. Gershman, S. J., & Blei, D. M. A tutorial on Bayesian nonparametric models. J. Math. Psychol. 56, 1-12 (2012).

40. Müller, P., & Quintana, F. A. Nonparametric Bayesian Data Analysis. Stat. Sci. 19, 95-110 (2004).

41. Speed, D., & Balding, D. J. MultiBLUP: improved SNP-based prediction for complex traits. Genome Res. 24, 1550-1557 (2014).

42. Lappalainen, T. et al. Transcriptome and genome sequencing uncovers functional variation in humans. Nature 501, 506-511 (2013).

43. 1000 Genomes Project Consortium. An integrated map of genetic variation from 1,092 human genomes. Nature 491, 56-65 (2012).

44. Zou, H., & Hastie, T. Regularization and variable selection via the Elastic Net. J. R. Stat. Soc. Ser. B. 67, 301-320 (2005).

45. Splansky, G. L. et al. The third generation cohort of the National Heart, Lung, and Blood Institute’s Framingham Heart Study: design, recruitment, and initial examination. Am. J. Epidemiol. 165, 1328-1335 (2007).

46. Hu, Z. L., Park, C. A., Wu, X. L., & Reecy, J. M. Animal QTLdb: an improved database tool for livestock animal QTL/association data dissemination in the post-genome era. Nucleic Acids Res. 41, D871-D879 (2013).

47. Spiliopoulou, A. et al. Genomic prediction of complex human traits: relatedness, trait architecture and predictive meta-models. Hum. Mol. Genet. 24, 4167-4182 (2015).

48. Goddard, M. E., Hayes, B. J., & Meuwissen, T. H. E. Using the genomic relationship matrix to predict the accuracy of genomic selection. J. Anim. Breed. Genet. 128, 409-421 (2011).

49. Lee, S. H., Weerasinghe, W. M. S. P., Wray, N. R., Goddard, M. E., & van der Werf, J. H. J. Using information of relatives in genomic prediction to apply effective stratified medicine. Scientific Reports 7, 42091 (2017).

50. Carbonetto, P., & Stephens, M. Scalable variational inference for Bayesian variable selection in regression, and its accuracy in genetic association studies. Bayesian. Anal. 7, 73-108 (2012).

51. Yi, H., Breheny, P., Imam, N., Liu, Y., & Hoeschele, I. Penalized multimarker vs. single-marker regression methods for genome-wide association studies of quantitative traits. Genetics 199, 205-222 (2015).

52. Sun, S. et al. Differential expression analysis for RNAseq using Poisson mixed models. Nucleic Acids Res. gkx204. doi: 210.1093/nar/gkx1204 (2017).

53. Tung, J., Zhou, X., Alberts, S. C., Stephens, M., & Gilad, Y. The genetic architecture of gene expression levels in wild baboons. Elife 4, e04729 (2015).

54. Zhou, X. et al. Epigenetic modifications are associated with inter-species gene expression variation in primates. Genome Biol. 15, 1 (2014).

55. Lea, A. J., Tung, J., & Zhou, X. A Flexible, Efficient Binomial Mixed Model for Identifying Differential DNA Methylation in Bisulfite Sequencing Data. PLoS Genet. 11, e1005650 (2015).

56. Manolio, T. et al. Finding the missing heritability of complex diseases. Nature 461, 747-753 (2009).

57. Shi, J. et al. Winner’s Curse Correction and Variable Thresholding Improve Performance of Polygenic Risk Modeling Based on Genome-Wide Association Study Summary-Level Data. PLoS Genet. 12, e1006493 (2016).

58. Li, J., Das, K., Fu, G., Li, R., & Wu, R. The Bayesian lasso for genome-wide association studies. Bioinformatics 27, 516-523 (2011).

59. Blei, D. M., & Jordan, M. I. Variational inference for Dirichlet process mixtures. Bayesian. Anal. 1, 121-143 (2006).

60. Blei, D. M., Kucukelbir, A., & McAuliffe, J. D. Variational inference: A review for statisticians. J. Am. Stat. Assoc. (in press), Preprint at https://arxiv.org/abs/1601.00670 (2017).

61. Ranganath, R., Tran, D., & Blei, D. M. (2016). *Hierarchical variational models*. Paper presented at the International Conference on Machine Learning.

62. Zhou, X. A Unified Framework for Variance Component Estimation with Summary Statistics in Genome-wide Association Studies. Ann. Appl. Stat. (in press), Preprint at http://biorxiv.org/content/early/2016/03/08/042846 (2017).

63. Andrews, D. F., & Mallows, C. L. Scale mixtures of normal distributions. J. R. Stat. Soc. Ser. B. 36, 99-102 (1974).

64. Verbyla, K. L., Hayes, B. J., Bowman, P. J., & Goddard, M. E. Accuracy of genomic selection using stochastic search variable selection in Australian Holstein Friesian dairy cattle. Genet. Res. 91, 307-311 (2009).

65. Wen, X., Luca, F., & Pique-Regi, R. Cross-Population Joint Analysis of eQTLs: Fine Mapping and Functional Annotation. PLoS Genet. 11, e1005176 (2015).

66. Harrow, J. et al. GENCODE: the reference human genome annotation for The ENCODE Project. Genome Res. 22, 1760-1774 (2012).

67. Stegle, O., Parts, L., Piipari, M., Winn, J., & Durbin, R. Using probabilistic estimation of expression residuals (PEER) to obtain increased power and interpretability of gene expression analyses. Nat. Protoc. 7, 500-507 (2012).

68. Guan, Y., & Stephens, M. Practical Issues in Imputation-Based Association Mapping. PLoS Genet. 4, e1000279 (2008).

69. Howie, B. N., Donnelly, P., & Marchini, J. A Flexible and Accurate Genotype Imputation Method for the Next Generation of Genome-Wide Association Studies. PLoS Genet. 5, e1000529 (2009).

70. Lee, S. H., Clark, S., & van der Werf, J. Estimation Of Genomic Prediction Accuracy From Reference Populations With Varying Degrees Of Relationship. bioRxiv, Preprint at http://biorxiv.org/content/early/2017/03/22/119164 (2017).

